# BPQ_RNA: A Thermodynamically Informed, Hybrid Quantum-Classical Framework for RNA 2D Structure Prediction Including Chemical Modifications

**DOI:** 10.64898/2026.11.25.769245

**Authors:** Nivedita Dutta, Liwen Shih, Quentin Vicens

## Abstract

Predicting RNA secondary (2D) structural folds remains a Nondeterministic Polynomial-time (NP)-hard challenge. Existing quantum approaches predominantly rely on coarse-grained heuristics or classical stem pre-estimation, shifting core thermodynamic computations outside the quantum workflow and limiting localized structural precision. Here, we present a Base-Pair QUBO framework for RNA (BPQ_RNA), a hybrid pipeline that directly maps experimental Nearest Neighbor Database (NNDB)-derived free energies onto a gate-based quantum framework strictly at the base-pair level. This eliminates pre-assembly constraints and integrates chemical modifications with zero additional qubit overhead. Executed on a GPU-accelerated simulator with classical maturation for multi-body loops, BPQ_RNA resolves biological ground states within a 5-22 qubit footprint. Finally, we analyze a Variational Convergence Bound, demonstrate how uniform randomized multi-start initialization and multi-replica sampling aids navigation of rugged energy landscapes, and outline future directions for scaling to more complex RNA 2D folds and expanded modification libraries.

## Introduction

RNA folding plays key roles in critical cellular functions, ranging from gene regulation to viral replication [1,2]. RNA folding involves the formation of Watson-Crick A-U and G-C base pairs, which together form the secondary structure of a particular RNA fold. Predicting the RNA secondary structure from sequence composition alone remains a foundational challenge in computational biology [3,4], due to shallow thermodynamic energy landscapes harboring competing local minima [3,5]. Hence, a single RNA sequence can adopt multiple secondary structures that exist in dynamic equilibrium [1]. This conformational equilibrium is frequently altered by cellular factors like post-transcriptional modifications or protein binding, which shift the energetic preference toward different structural states.

Given the highly rugged and non-convex nature of this energetic landscape, quantum optimization has emerged as a promising computational alternative capable of tunneling through deep local minima [6]. Prior formulations mapped structural configurations to QUBO equations using either coarse topological heuristics over classically pre-assembled macroscopic stems [7,8] or stacked quartets of length two [9]. These foundational models successfully demonstrated proof-of-concept 2D structural predictions for short RNA transcripts, establishing the applicability of quantum algorithms in computational RNA biology. However, macroscopic stem architectures introduce severe qubit overheads and steep energy cliffs, while quartet-level variables rely on unphysical, arbitrarily tuned soft constraints that mask localized nucleotide variances.

Modern utility-scale implementations suffer from unconditioned energy landscapes that necessitate manual tuning of the underlying quantum circuit architecture (the “ansatz”) to converge. Recent benchmarks by Andrews et al. (2026) highlighted the “missing physics” in standard QUBO models: restricting the optimization domain strictly to two-body interactions fundamentally fails to capture the non-local energetic consequences of multi-body loops [10]. This structural limitation creates deep local minima that trap variational solvers, frequently leading to the severe over- or under-prediction of key interactions like pseudoknots [10]. Resolving this thermodynamic gap necessitates an integrative hybrid approach that can map exact physical parameters while resolving complex energetics.

Recent advancement in quantum-centric combinatorial optimization of RNA secondary structures has enabled successful execution of QUBO models on physical quantum hardware using up to 156 qubits, utilizing sophisticated error mitigation and heuristic sampling schemes [11]. In this framework, decision variables are defined over pre-assembled quartets (stacked 2-base-pair units), each assigned an empirical free energy. Extending stems across adjacent quartets then requires adding auxiliary reward multipliers and soft boundary penalties [11]. While this demonstrates impressive algorithmic scaling, particularly in the approach reported by Kumar and coworkers [11] towards modeling complex non-planar interactions like pseudoknots, it does not map individual base-pair stacking free energies directly into the quadratic Hamiltonian. As recent evaluations have highlighted, expanding the topological search space with uncalibrated energy landscapes frequently leads to the overestimation of motifs and steep drop-offs in predictive accuracy [10]. This underscores that hardware scalability alone is insufficient; the underlying quantum Hamiltonian must be intrinsically grounded in exact biophysical parameters.

To address these architectural limitations, we introduce a localized, base-pair level hybrid quantum-classical pipeline. By mapping binary quantum variables strictly onto individual base pairs, adjacent nearest-neighbor stacking rewards (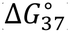) accumulate via quadratic interactions in the quantum Hamiltonian. We apply a microscopic linear bias directly into the cost Hamiltonian to flatten rugged energy valleys, guiding the classical optimizer to approach the global minima. This eliminates classical stem pre-assembly and allows the efficient integration of RNA modifications with zero additional qubit overhead. We also incorporate N6-methyladenosine (*m*^6^*A*), one of the most abundant RNA posttranscriptional modifications that plays crucial roles in regulating RNA stability, gene regulation and cellular homeostasis, while driving diverse pathological conditions including cancer progression [12]. By updating the underlying thermodynamic parameters rather than expanding the variable space, our framework seamlessly integrates *m*^6^*A* within a compact quantum workflow.

The present study serves as an exploratory proof-of-concept to examine the feasibility of predicting stem-loop conformations using a base-pair level QUBO designed for modest, widely accessible computational environments. Non-local structural features, including size-dependent hairpin loops and asymmetric dangling ends, are subsequently handled via a classical maturation layer post-collapse to avoid higher-order hardware constraints. Executed on a statevector simulator utilizing standard single-GPU acceleration (e.g., NVIDIA T4) within a 5-22 qubit envelope, our pipeline combines random multi-start and multi-replica sampling with generative pooling to generate an energetic ranking of candidate conformers across independent optimization trajectories. This allows the pipeline to aggregate diverse set of conformers, yielding a 2D structural ensemble of sampled RNA folds.

## Methods

### Theoretical Framework and Base-Pair Level Mapping

We establish our optimization domain over a vector of binary variables *x*_*u*_ ∈ {0, 1}, where *u* = *(i, j)* represents the physical formation of a single discrete intramolecular base pair between the *i*-th and *j*-th nucleotides of an RNA sequence. Candidate variables are instantiated if and only if they satisfy pre-defined pairing rules (canonical A-U, U-A, C-G, G-C; wobble G-U, U-G; and modified M-U, U-M) alongside a strict steric constraint enforcing a minimum loop separation of *j* ≥ *i* + 4. The global mathematical optimization objective is initially formulated as a classical QUBO objective *O(x)* to be minimized:

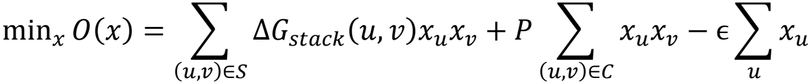

Here, *S* denotes valid continuous stems (where *u* = (*i, j*) and *v* = (*i* + 1, *j* − 1)), with stabilizing weights *ΔG*_stack_(*u, v*) extracted directly from empirical NNDB tables (e.g., −34 *kcal/mol* for a 5’-GC-3’ / 3’-CG-5’ stack) [12-14]. As these free energy values are stabilizing (negative), stem elongation is driven by the quantum solver minimizing the objective function (i.e., maximizing the magnitude of these terms), favoring conformations with lower total stacking free energy. The structural conflict set *C* penalizes base sharing (*i* = *k, j* = *l*, etc.) or crossing base pairs (*i* < *k* < *j* < *l*), corresponding to non-planar pseudoknots using a *LinearInequalityToPenalty* transformation with a fixed constraint penalty multiplier P = 50.0 kcal/mol. Consequently, while the Hamiltonian utilizes empirical free energies, the inclusion of constraint penalties renders the QUBO cost landscape distinct from the thermodynamic free energy landscape of RNA. However, fixing the penalty multiplier (*P* ≫ |Δ*G*_stack_|) strictly prevents unfeasible structures without inducing optimization instabilities from auto-scaling penalties. In this framework, the structural domain is restricted to planar secondary structures, explicitly excluding pseudoknots. As recently highlighted by Andrews et al. [10], mapping the context-dependent energetics of pseudoknots onto a second-degree polynomial (QUBO) requires introducing heuristic penalties that frequently result in severe structural overestimation. To ensure that our native NNDB thermodynamic mapping is evaluated strictly on its physical fidelity without the noise of arbitrary topological tuning, pseudoknot formation is reserved for future higher-order implementations.

To mitigate barren plateaus, a microscopic symmetry-breaking bias (ϵ = −0.01) slopes the cost landscape into guided valleys without altering relative thermodynamic rankings. This base-pair resolution integrates post-transcriptional modifications (*e. g. m*^6^A), with zero qubit overhead. By treating *m*^6^A as a native character (‘M’) in the sequence alphabet, candidate generation seamlessly maps (M-U) and (U-M) variables. Incorporating its updated thermodynamic stability [15] requires only updating the *ΔG*_stack_ dictionary lookup matrix without modifying the underlying structural constraint equations or expanding the qubit register.

To solve the resulting QUBO model, our approach relies on a Variational Quantum Algorithm (VQA), a hybrid approach that pairs a parameterized quantum circuit with a classical optimizer [16]. Specifically, we use the Quantum Approximate Optimization Algorithm (QAOA) [17], which is tailored for combinatorial problems. To apply the QAOA, the variable space is mapped onto quantum spins via standard algebraic transformation: 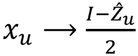, where 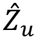 is the Pauli-Z operator acting on the physical qubit mapped bijectively to candidate base pair *u*, and *I* is the identity operator. Substituting this mapping into the QUBO objective function yields the final problem Ising Hamiltonian *H*_P_:

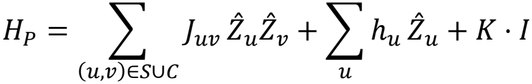

### Hybrid Pipeline Architecture and GPU-Accelerated Execution

Our architecture decouples two-body and multi-body thermodynamics. Candidate pairs (*j* >= *i* + 4) are compiled into a QUBO and optimized via a multi-cycle QAOA circuit (4 layers, COBYLA optimizer, 500 max iterations) simulated on a cloud-provisioned NVIDIA T4 GPU (Google Colaboratory) via qiskit-aer [18].

Upon quantum statevector collapse and sampling, the optimal statevector is routed to a classical maturation layer that applies non-local multi-body corrections utilizing empirical NNDB parameters: size-dependent hairpin destabilizations at the innermost boundary (min(*j* − *i*)) and asymmetric dangling end stabilizations at the outermost boundary (max(*j* − *i*)), The total free energy is output as:

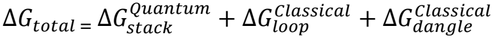

Where 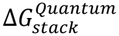 is the continuous stabilizing stacking energy evaluated from the QAOA-optimized QUBO formulation, 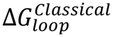 is the classical energetic correction for size-dependent hairpin loops and 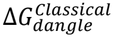 is the correction for unpaired terminal nucleotides.

### Multi-Replica, Multi-Start Protocol with Dynamic Early Stopping

To evaluate solver stability and sample sub-optimal conformational landscapes, execution was evaluated across the 20 RNA test sequences (*S*1-*S*20, sequence lengths 10-16 nt), generating QUBO formulations requiring up to 22 qubits. Each sequence was processed across 5 independent solver replicas. To navigate non-convex energy landscapes in dense conflict graphs without incurring redundant compute, each replica implemented a uniform randomized multi-start initialization scheme capped at a maximum of *N* = 20 variational parameter cycles (*γ, β*) using QAOA (*number of circuit layers, p* = 4) and a classical COBYLA optimizer (maximum 500 iterations). An adaptive early-stopping criterion terminates the cycle loop immediately upon resolving the exact classical QUBO ground state. While Python-level preprocessing is secured with a global random seed, the GPU-accelerated QAOA solver is unconstrained to maximize exploratory sampling across replicates.

Upon completion of the optimization routine, candidate bitstrings were extracted from the final measured probability distribution generated by Qiskit BackendSamplerV2 *primitive* [18]. The top *K* = 5 most frequently sampled unique bitstrings per replica were retained and passed to the classical maturation layer to evaluate complete thermodynamic parameters (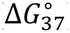). This yielded a benchmark dataset of 500 fully evaluated candidate folds (20 sequences × 5 replicas × 5 ranks).

## Results and Discussion

To evaluate the predictive performance and computational efficiency of the BPQ_RNA framework, we benchmarked the pipeline on a dataset of 20 stem-loop/hairpin motifs (10-16 nucleotides, sequences S1-S20; Table 1). This benchmark dataset encompasses both unmodified canonical sequences and *N*^6^-methyladenosine (*m*^6^A)-modified variants, including biologically relevant constructs derived from the human *Xist* RNA A-repeat and tRNA anticodon stem-loops. For these test sequences, we have quantified the candidate(base-pair)-density qubit register scaling, evaluated variational convergence dynamics across rugged constraint landscapes, and benchmarked 2D secondary structure predictions against experimental ground states and classical predictive baselines (MXfold2 [20] and RNAstructure [21]). To explore a thermodynamically grounded quantum framework, our pipeline serves as a small-scale proof-of-concept that directly maps empirical NNDB-derived stacking free energies (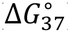) into the quadratic terms of the Hamiltonian (Fig.1). Rather than attempting to force complex multi-body physics into a purely quadratic quantum framework, which often introduces unconditioned energy cliffs [10], we evaluate a pragmatic division of tasks. The quantum core is tasked solely with native two-body base-pairing interactions, while non-local loop energetics are handled by a classical maturation layer post-collapse.

**Fig. 1.**
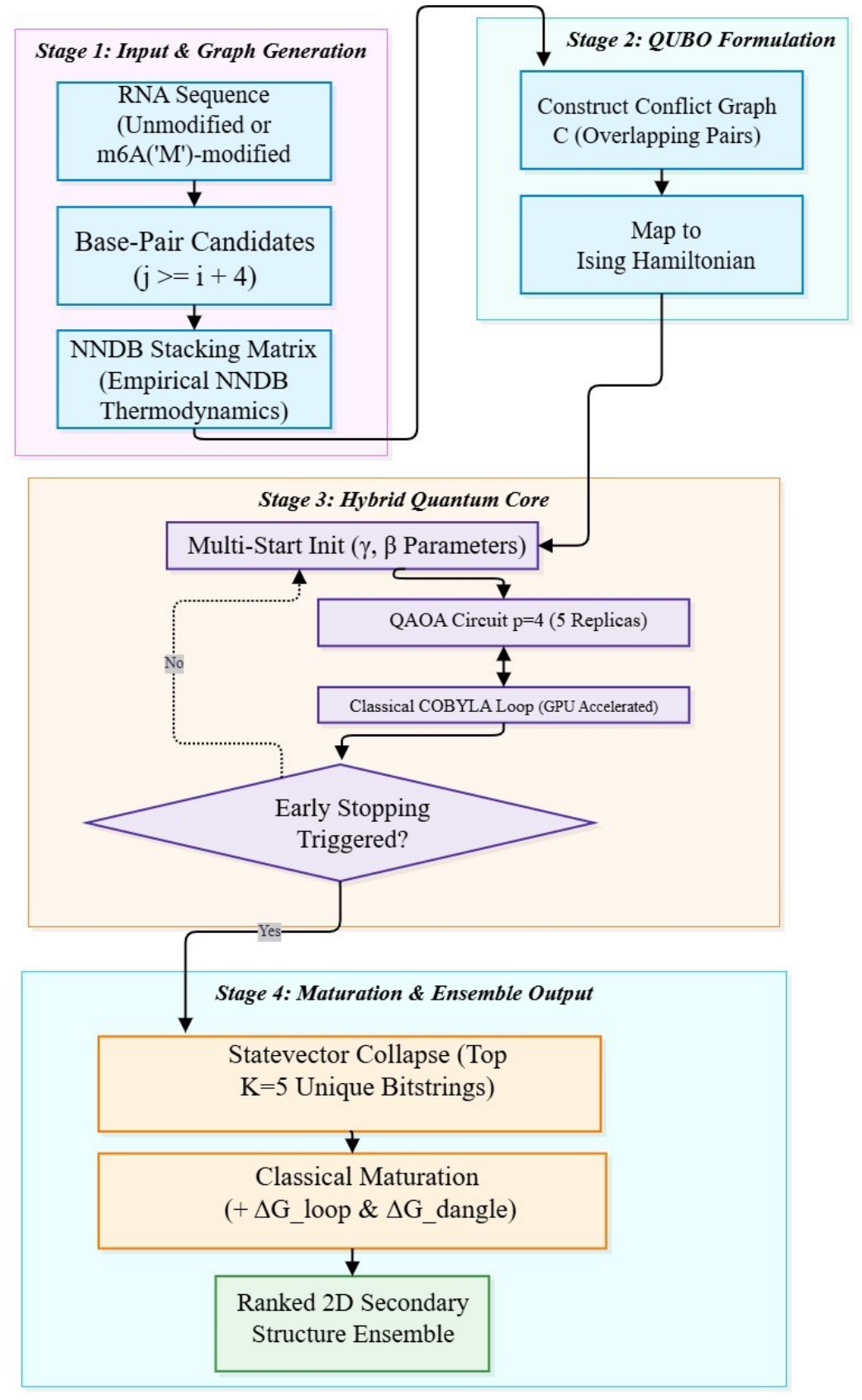
BPQ_RNA hybrid quantum-classical RNA 2D-structure prediction workflow. (A) Overview of the four-stage BPQ_RNA framework. *Stage 1* compiles candidate base pairs, dynamically embedding empirical NNDB base stacking free energies, including the incorporation of *m*^6^A modifications; *Stage 2* strictly maps these two-body interactions and structural constraints to an Ising Hamiltonian; *Stage 3* highlights the GPU-accelerated quantum core, utilizing a randomized multi-start QAOA (*γ, β* parameters, *p* = 4 layers) with adaptive early stopping for computational efficiency; and *Stage 4* executes classical maturation on the top K=5 sampled bitstrings to resolve the multi-body loop and dangling end energetics (obtained from NNDB parameters), extracting a thermodynamically ranked 2D structural ensemble directly from the probabilistic quantum measurement profile. The flowchart was generated using draw.io Desktop, version 32.5.5 [19].

**Table 1.**
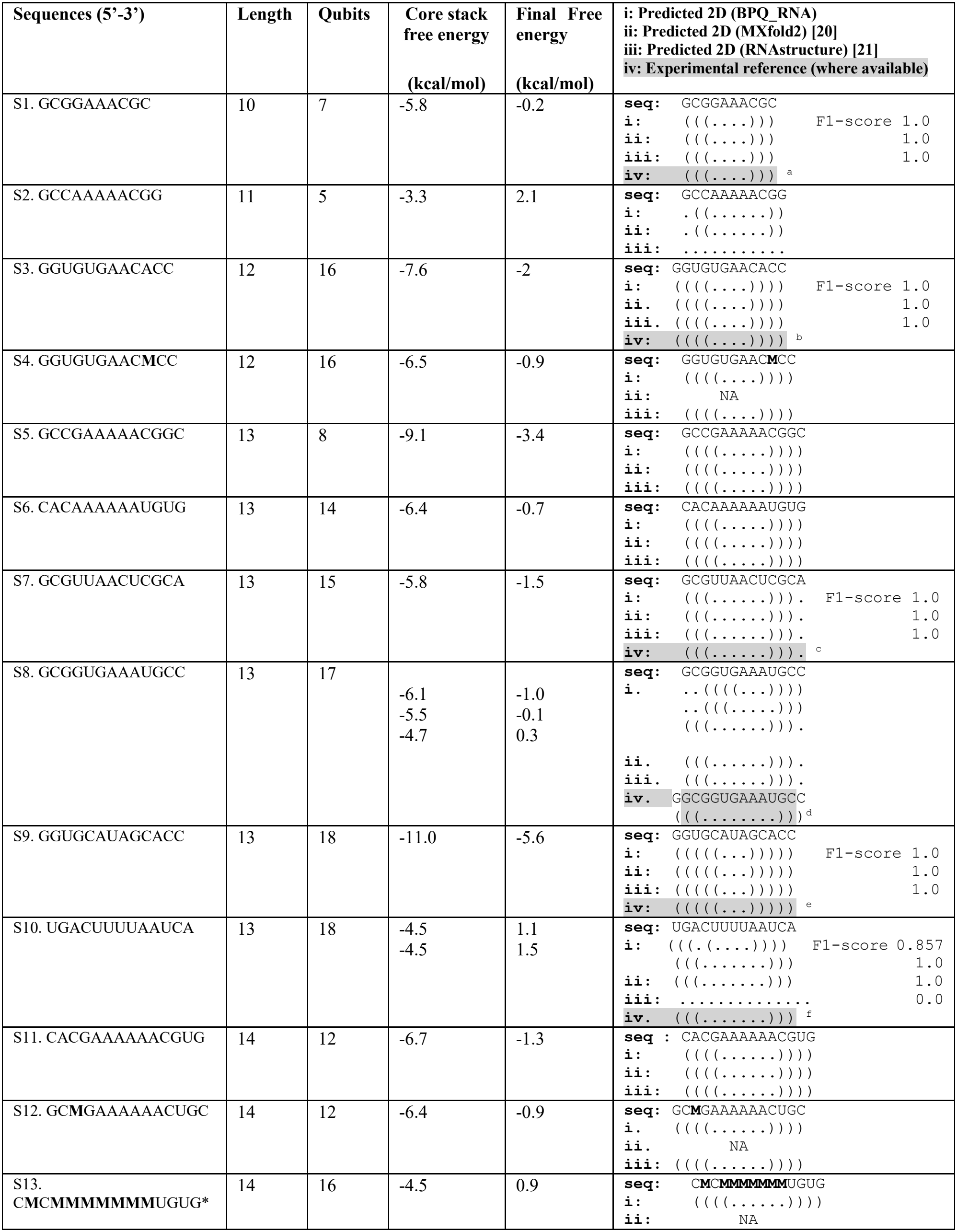
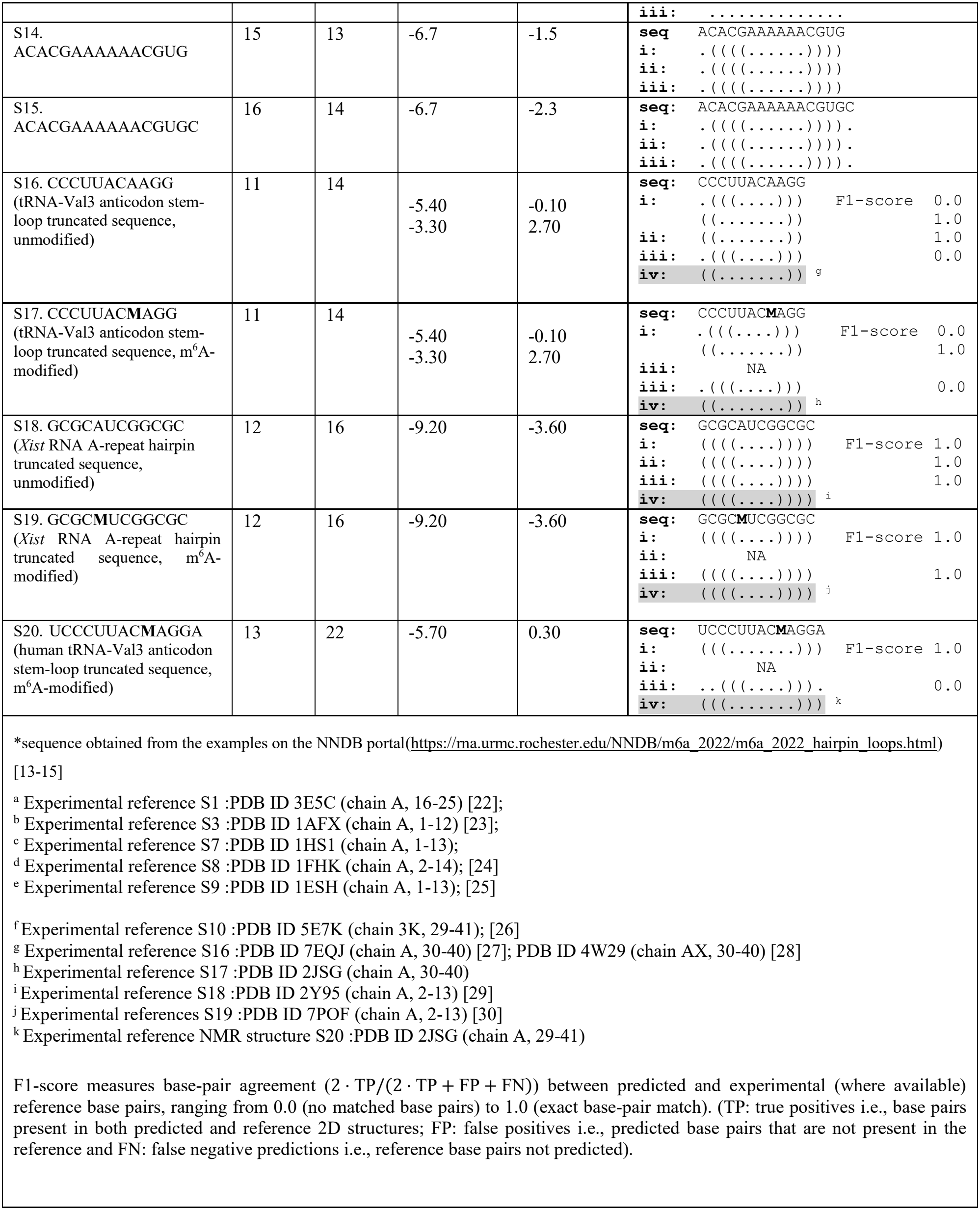
Predicted 2D structures and associated energetics for the 20 test sequences.

### Benchmark of BPQ_RNA on unmodified and modified stem-loop motifs

In this work, our benchmark focuses on small-scale predictive analysis for stem-loop (hairpin) motifs (with or without dangling ends). By mapping decision variables strictly at the individual base-pair level (1 qubit per candidate pair), our hybrid base-pair QUBO formulation was able to reproduce the experimental 2D structure (where available) or make predictions in close agreement with the other predictors, within a compact 5-22 qubit footprint for the test sequences (**Table 1; Fig 2A**). As illustrated in **Fig.2A**, the required qubit register size is intrinsically driven by the base-pairing density of the specific sequence context. Since each qubit represents one candidate base pair, the resource density can be quantified as candidate base pairs per nucleotide (Qubits/nt). For example, comparing S1 (10 nt, 7 qubits) and S2 (11 nt, 5 qubits) demonstrates that a longer sequence can require a smaller qubit register if its candidate pairing space is structurally constrained. Similarly, among 13-nucleotide sequences, the structural sparsity of S5 requires only 8 qubits (~0.6 Qubits/nt), whereas the increasing self-complementarity of S7 (15 qubits, ~1.2 Qubits/nt), S9 (18 qubits, ~1.4 Qubits/nt), and S20 (22 qubits, ~1.7 Qubits/nt) progressively expands the qubit register. m^6^A-modified sequences (marked with asterisks) map to the exact same qubit count as their unmodified counterparts (e.g., S3 vs. S4; S18 vs. S19), reinforcing the zero-overhead advantage of updating local thermodynamic matrices.

**Fig. 2.**
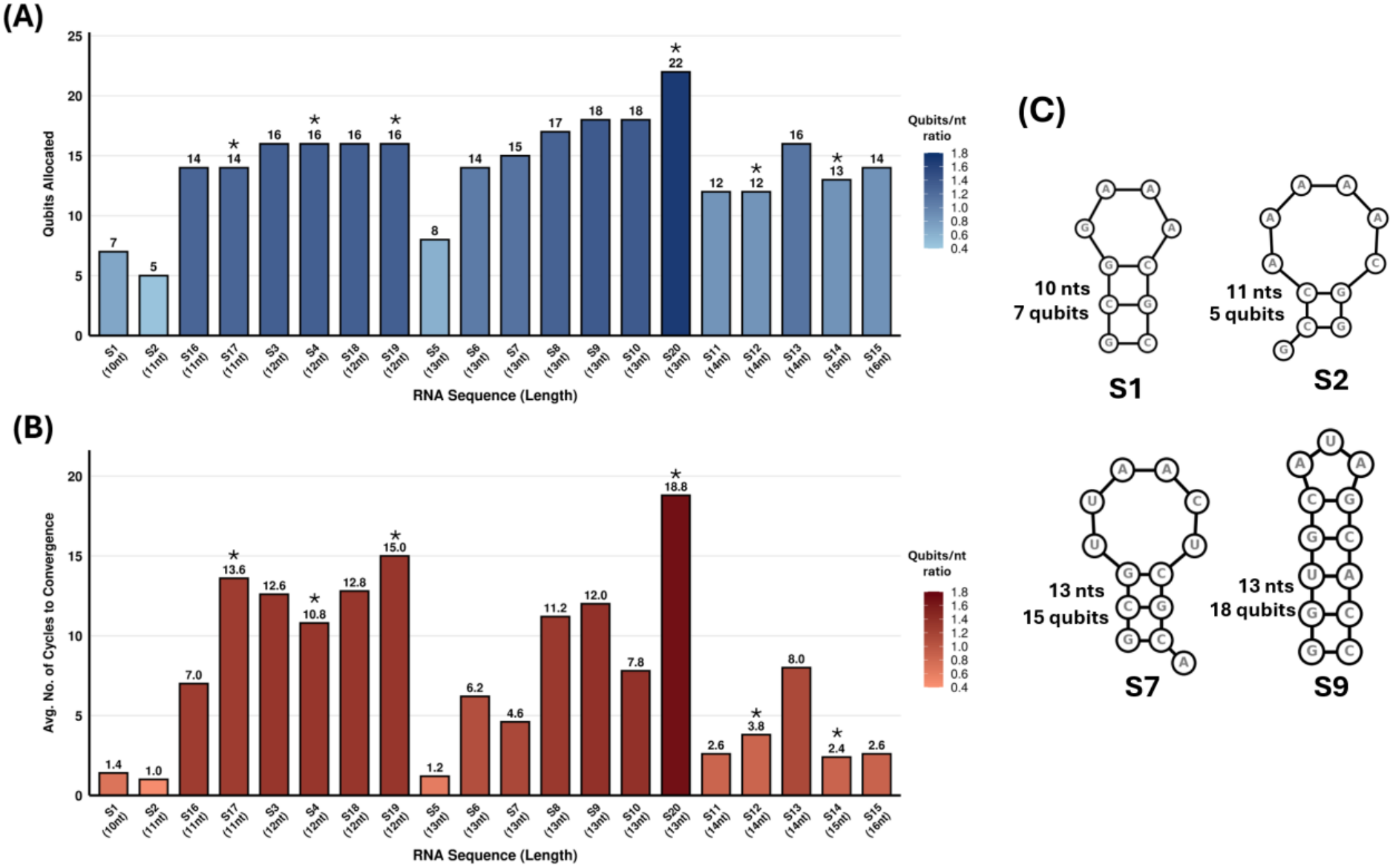
Quantum resource scaling and variational convergence dynamics. (A) *Qubit allocation for test sequences S1-S20*: The *m*^6^A-modified sequences are indicated with asterisks. The varying sequence contexts influence the number of qubits allocated for sequences of the same length. (B) *Average number of randomized multi-start cycles required to convergence*: The number of allocated qubits and the sequence context for the test sequences were observed to impact average number of cycles required to converge to the global core stack energy minimum. The interplay between qubit-to-nucleotide ratio and the ruggedness of the constraint landscape due to the sequence context, dictates the necessary number of adaptive multi-start initializations required to avoid local traps and resolve the global core stacking free energy minimum. (C) BPQ_RNA predicted 2D structures for representative test sequences S1, S2, S7, and S9: S1 (10 nts, 7 qubits) and S2 (11 nts, 5 qubits); while S7 (13 nts, 15 qubits) and S9 (13 nts, 18 qubits). This illustrates qubit register allocation scaling depends on the base-pairing density and sequence context.

Benchmarking BPQ_RNA against established computational tools reveals that our hybrid formulation consistently matches the predictive accuracy of legacy thermodynamic algorithm (RNAstructure) and thermodynamic integration combined with deep learning (MXfold2), as detailed in Table 1. For the unmodified RNA sequences, BPQ_RNA accurately evaluates the overall thermodynamic stability by accounting for Watson-Crick stacking, loop and dangle energetics. In standard short-stem/short-loop motifs such as S1, S3 and S9, BPQ_RNA converges precisely on the expected 2D structures respectively, matching both the experimental reference 2D structures and classical solver outputs. For S4, S5, S6, S11, S14 and S15, for which the experimental 2D is not known, the BPQ_RNA predictions are observed to agree with MXfold2 and RNAstructure predictions. As sequence complexity increases to include larger asymmetric loops and dangling ends (e.g., S9, S14, and S15), BPQ_RNA correctly allocates qubits to the continuous native stems (yielding strong quantum core stacking free energies) while classically resolving the effect of the terminal un-paired nucleotides. For S8, while a 5’-terminal truncation in the test sequence prevents a strict one-to-one F1-score calculation against the experimental reference, BPQ_RNA’s multi-replica sampling successfully captures the same sub-optimal stem-loop topology “(((…..))).” (containing a dangling 3’C) predicted by MXfold2 and RNAstructure. Three of the sampled conformers for S8 from the pool are shown in Table 1, to showcase the alternate predictions by BPQ_RNA.

### Predicting 2D structure of RNA sequences with *m*^6^*A* Modifications

From a structural biology perspective, the integration of chemical and post-transcriptional modifications, specifically *N*^6^-methyladenosine (*m*^6^*A*) (**Fig.3A**), provides a substantial architectural advantage over existing quantum-centric approaches which don’t take modified residues into account. Prior QUBO formulations [6-11] have exclusively restricted the RNA alphabets to standard canonical and wobble base pairing within unmodified bases (A, U, G and C), entirely overlooking the thermodynamic consequences of post-transcriptional modifications. As highlighted in a recent review by Yang et al. (2025) [31], despite rapid advancements in classical physics, statistics and deep learning-based architectures for RNA secondary structure prediction, accurately modeling RNA modifications remains a critical challenge. From a structural biology perspective, our BPQ_RNA framework directly addresses this challenge by integrating post-transcriptional modifications, specifically *N*^6^-methyladenosine (m^6^A), into a gate-based quantum workflow with zero additional qubit overhead. Here, we selected m^6^A as it is the most abundant internal post-transcriptional modification in eukaryotic mRNA, while also frequently occurring in other types of RNA (such as tRNA), playing essential roles in RNA stability, splicing, and translation, while driving diverse pathological conditions [12, 31].

The BPQ_RNA framework treats modified residues directly at the continuous stacking level by updating the underlying Nearest Neighbor Database (NNDB) energy matrices (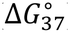). Because *m*^6^*A* alters nearest-neighbor stacking free energies (Fig. 3B) without expanding the candidate base-pair graph, replacing Adenine with *m*^6^*A* (‘M’) adjusts the quadratic coupling constants (*J*_*uv*_) in the Ising Hamiltonian without increasing the required qubit register relative to the unmodified sequence.

**Fig. 3.**
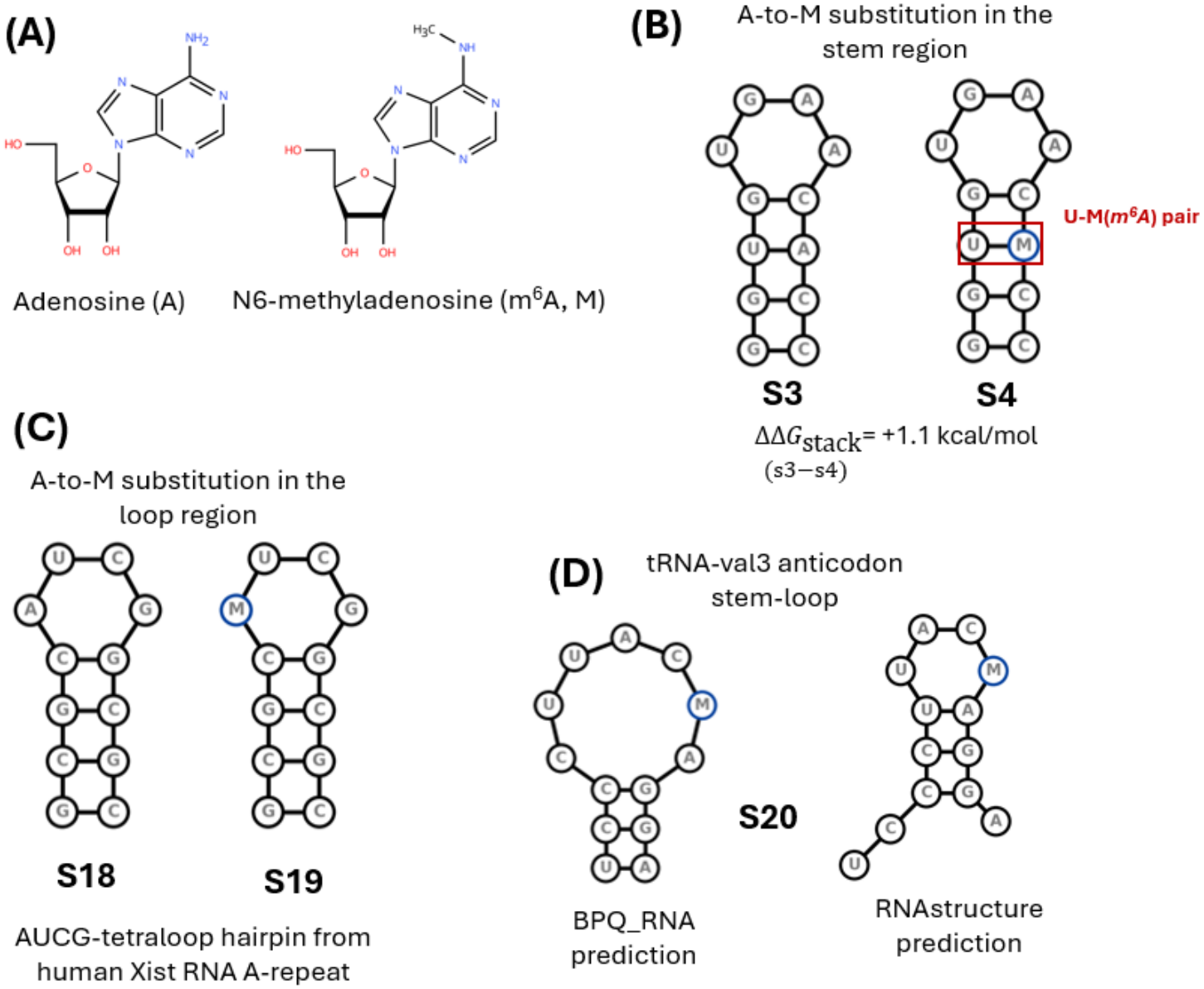
2D structural prediction of *m*^6^A-modified RNA motifs by BPQ_RNA. **(A)** Chemical structures of Adenosine (A) and *N*^6^-methyladenosine (*m*^6^A, M). **(B)** Helical destabilization induced by an *m*^6^A ⋅ U pair in stem sequence **S4** relative to unmodified **S3** (ΔΔ*G*_stack_ = +1,1 kcal/mol). **(C)** Positional accommodation of *m*^6^A within the single-stranded loop of human *Xist* RNA A-repeat hairpin (**S18, S19**). **(D)** Comparison of the human tRNA-Val3 anticodon stem-loop (**S20**) 2D structure predicted by BPQ_RNA versus classical RNAstructure [21] tool, demonstrating the sensitivity of the BPQ_RNA hybrid quantum-classical prediction framework to chemical modification energetics.

To validate this, we tested the unmodified sequences and their *m*^6^A-modified (‘M’) counterparts, including experimentally determined biologically relevant stem-loop motifs. Comparing S3 to S4 and S18 to S19 demonstrates that BPQ_RNA correctly preserves the structural integrity of these robust “((((……))))” stem-loops, identically matching the MFE (minimum free energy) predictions by RNAstructure (Table S1, Fig. 3C). The core stack free energy and final free energy values obtained using BPQ_RNA, reflect the thermodynamic destabilization by *m*^6^A modification in the stem for S4 compared to S3 (Fig. 3B).

For S12, we tested another example with *m*^6^A in the stem and BPQ_RNA predicted the 2D structure in agreement with the RNAstructure prediction. For sequences S18 and S19, the free energy values for the unmodified and modified 2D structures were predicted to be the same, suggesting *m*^6^A in the loop region doesn’t alter the structure at the 2D level for this sequence as supported by the experimental (NMR) 3D-derived 2D structures for this AUCG-tetraloop hairpin from human Xist RNA A-repeat [29,30]. Similarly, for the sequence constructs derived from the *E*.*coli* tRNA-val3 anticodon stem-loop sequence [27,28] (unmodified (S16) and *m*^6^A-modified (S17)), are predicted to have the same 2D structures. For these sequences, while the MFE structure obtained from BPQ_RNA and RNAstructure [21] differs from the experimental structure (F1-score 0.0), BPQ_RNA also sampled the alternative lower stability conformer for both sequences in agreement (F1-score 1.00) with the experimental observation. MXfold2 [20] which although doesn’t predict modified RNA 2D structure, accurately predicted the 2D for the unmodified sequence S16.

For the longer construct S20 (also derived from tRNA-val3 anticodon stem-loop containing *m*^6^A37), from our test run (5 replicas), the BPQ_RNA predicted 2D structure was observed to be in agreement with the experimental observation while RNAstructure prediction was found to have a F1-score of 0.0 (Fig. 3D). For a highly modified sequence (saturated with a continuous *m*^6^A tract (CMCMMMMMMMUGUG), while BPQ_RNA predicted “((((……))))” stem-loop structure, the final free energy (+ve) of 0.90 showcases the instability of this modified motif. In contrast, RNAstructure predicted a completely unpaired “…………..” structure which it does for any sequence for which the MFE value is calculated to have a positive value. For example, both BPQ_RNA and MXfold2 predicted a stem-loop motif with a dangling G-nucleotide for S2 and a stem-loop with a 3-base-pair stem for S10 ; (RNAstructure predicted an unpaired 2D structure for both). For S10 (sequence corresponding to the tRNALys anticodon stem-loop [26], the comparison with the experimental 2D structure demonstrates that BPQ_RNA and MXfold2 predictions agree with the experimental structure.

Our benchmark comprising 20 test sequences corresponding to such a simple motif (stem-loop) is able to showcase the differences between the physics-based RNAstructure and integrative physics and deep-learning based MXfold2, while demonstrating the applicability of random-start, multi-replica approach for the prediction/sampling of RNA 2D structural ensembles for higher probability to detect the alternative 2D fold especially for chemically modified RNA sequences. These results suggest that RNA modifications can be incorporated into gate-based quantum optimization purely through energetic parameterization, establishing a proof-of-concept blueprint for expanding quantum workflows to broader modification libraries (e.g., pseudouridine, inosine).

### Variational Convergence Bounds and Optimization Dynamics

Our benchmark reveals the algorithmic behavior of the BPQ_RNA framework on dense candidate graphs, specifically highlighting how penalty scaling shapes the variational landscape. In sequence regions featuring highly overlapping complementary interactions or dense self-complementarity, such as the complex folding landscape of S20 (UCCCUUACMAGGA, requiring 22 qubits for just 13 nt), the densely paired S9 (GGUGCAUAGCACC, 18 qubits for 13 nt), or the repetitive GC-rich stretch in S18 (GCGCAUCGGCGC, 16 qubits for 12 nt), candidate edge hyper-saturation creates severely rugged optimization landscapes dominated by steep penalty cliffs (Table S1; Fig. 2B).

Specifically, highly complementary or GC-dense sequences allow individual nucleotides to form valid Watson-Crick or Wobble base pairs with multiple alternative partners across the RNA chain. In a QUBO formulation where binary variables (*x*_*u*_ ∈ {0,1}) represent the formation of individual base pairs, this combinatorial multiplicity drastically inflates the algorithmic variable density (qubit-to-nucleotide ratio). Because the current approximation of the framework restricts a nucleotide to participate in only a single base-pairing interaction at a time, all mutually exclusive candidate base pairs must be enforced via heavy quadratic penalty terms (P ⋅ *x*_*u*_*x*_*v*_, where P ≫ *l*Δ*G*_stack_*l*). When evaluating parameterized quantum states *lψ*(*γ, β*)⟩, statevectors that violate these dense non-overlap constraints, encounter steep penalty walls, driving intermediate objective function values into unfeasible, large positive raw objective values during intermediate QAOA cycles (ranging from +49.98 up to +2847.48). Under standard single-trajectory variational optimization, classical optimizers such as COBYLA become trapped behind these steep penalty cliffs in local minima corresponding to misfolded or incomplete pairing configurations. Dynamically scaling independent random-start initialization cycles (*γ, β*) enables QAOA to tunnel through these unfeasible constraint traps and converge reliably on the Minimum Free Energy (MFE) state.

The hyper-saturation trapping is directly observed in our benchmarking results for sequences with high qubit-to-length ratios. S18: Replicates 1 and 5 were trapped in local QUBO minima (yielding core stacking energies -5.84 kcal/mol and -5.83 kcal/mol, respectively) before completing all 20 optimization cycles, failing to discover the true QUBO global minimum of -9.24 kcal/mol; S20: Replicate 3 encountered severe penalty trapping in unfeasible regions (yielding an intermediate objective of +49.56 while Replicates 1, 2, and 5 also converged on suboptimal local minima (yielding core stacking energies of -4.54 kcal/mol to -0.03 kcal/mol). Only through dynamic multi-start initialization cycles was the solver able to tunnel through these unfeasible constraint barriers, enabling Replicate 4 of S20 to reach the exact global QUBO minimum (core stacking energy of - 5.74 kcal/mol) and correctly recover the MFE structure with 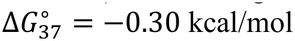). On the contrary, sparse motifs with limited valid pairings, such as the homopolymeric poly-A tracts in S6 and S11, maintain low variable density (0.86 to 1.08 qubits/nt) and routinely converge on the global minimum within 1 to 2 variational cycles without penalty trapping. Because escaping these dense penalty traps relies on unconstrained stochastic sampling, intermediate solver trajectories will naturally vary across independent executions. To ensure baseline reproducibility despite this probabilistic variance, the complete benchmark outputs for all 20 sequences are provided in **Supplementary Data 1**, allowing end-users to independently scale multi-start cycles and replicates based on available hardware.

While our hybrid workflow successfully identifies the experimental ground states for the considered sequences (where directly comparable), it explicitly navigates a fundamental limitation of quadratic formulations. Because the quantum objective function is inherently limited to second-degree polynomials (QUBO), it greedily optimizes for two-body stacking rewards 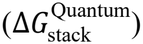. Macroscopic multi-body features, such as size-dependent hairpin loop destabilizations, internal loop asymmetries, and non-local topological penalties cannot be mapped into a strictly quadratic Hamiltonian without introducing heuristic penalty noise or resorting to highly resource-intensive higher-order polynomial unconstrained binary optimization (HUBO).

By decoupling these energetic contributions and deferring multi-body structural evaluations to the classical maturation layer post-collapse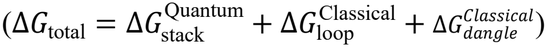, BPQ_RNA maintains close thermodynamic consistency with empirical free-energy-parameter-based RNA 2D structure prediction while operating within a compact 5-to-22-qubit simulation window. This architectural fragmentation highlights the broader computational challenge of fully quantum RNA folding, providing a pathway for future iterative CPU-QPU feedback mechanisms, where classical loop evaluations could dynamically update quantum stacking weights across multiple variational cycles. Overall, this modularity positions BPQ_RNA as a prospective subroutine within larger workflows. Rather than attempting to fold massive, full-length RNA sequences monolithically on a QPU, this local motif-focused approach can be selectively deployed to resolve and refine specific structural motifs.

### Adaptive Multi-start Convergence and Multi-Replica Sampling Across S1-S20

The integrated adaptive multi-start protocol demonstrated computational efficiency, effectively balancing rapid convergence with necessary energy landscape exploration. Across the 100 replica executions (20 sequences × 5 independent solver replicas), the early-stopping criterion successfully triggered in 88% of all runs (88/100) (Table S1). Short or structurally sparse sequences (e.g., S1, S2, S5, and S11) routinely converged on the Minimum Free Energy (MFE) structure almost immediately, averaging just 1.0 to 2.6 variational cycles and successfully avoiding unnecessary simulation overhead. On the other hand, sequences featuring dense conflict graphs and overlapping complementary regions (e.g., S9, S18, and S20) required extended multi-start sampling, averaging 12.0 to 15.0 cycles, to escape local energy traps, occasionally utilizing the full 20-cycle allowance to reach global convergence.

Across the 100 Rank-1 candidate evaluations, the pipeline recovered the exact classical QUBO ground state in 89 out of 100 individual instances (89%). Relying on a multi-replica ensemble proved highly robust against the isolated trapping events observed in denser optimization landscapes. By aggregating results across 5 independent initializations, the BPQ_RNA pipeline ultimately recovered the global MFE state for all of the test sequences, with 19 of those sequences identifying the ground state in at least a majority (3 out of 5) of their respective runs. Additionally, retaining the top K=5 unique bitstrings from the final measurement distribution allowed the classical maturation step to evaluate nearby low-energy configurations without requiring separate algorithmic executions. This post-collapse sorting successfully captured relevant sub-optimal structures in the surrounding conformational landscape, effectively generating an energetic ensemble of alternative 2D folds. In a biological context, capturing this distribution directly from the quantum measurement profile is highly advantageous for investigating RNA conformational equilibria.

## Discussion

Accurate prediction of RNA secondary structure remains difficult when accounting for the localized thermodynamic effects of chemical modifications and non-canonical base pairing interactions. Prior QUBO models [8-10] for RNA 2D structure optimizations heavily relied on stem-based encodings, where the fundamental binary variables represented pre-assembled macroscopic stems. In such formulations, modulating structural behaviour required applying heuristic one-body penalties to entire stems based on length or compositions, a strategy previously shown to fail at capturing emergent energetic correlations or preventing topological overestimation [10]. By operating strictly at the base-pair level, our BPQ_RNA framework circumvents this limitation. Within BPQ_RNA, stems are not rigidly pre-defined; rather they emerge dynamically through two-body interactions between adjacent base pairs. This allowed us to directly embed experimental NNDB stacking free energies into the quadratic terms of the Hamiltonian, replacing unphysical one-body heuristics with exact, correlated thermodynamics.

While recent large-scale quantum implementations have explored mRNA secondary structure prediction using advanced variational algorithms and tensor network simulations [11, 32], these approaches highlight a critical bottleneck: optimizing an unconditioned, heuristically derived energy landscape scales poorly in biological accuracy, regardless of qubit count or error mitigation. By contrast, our BPQ_RNA framework prioritizes physical fidelity over raw hardware scaling, by tasking the quantum core solely with native two-body base-pairing interactions, while non-local loop energetics are handled by a classical maturation/post-processing layer.

Capturing the 2D structural possibilities directly from the quantum measurement profile addresses a major limitation of purely thermodynamics-based or static deep-learning based predictions. Even for short, localized stem-loop configurations, state-of-the-art classical predictive tools (such as RNAStructure) can struggle to predict the accurate 2D structure if it resides in a slightly sub-optimal energetic basin, as we observe for S10, S16, S17 and S20. Therefore, the ability to efficiently generate a 2D structural ensemble, utilizing the inherent probabilistic sampling of quantum measurements to capture functionally critical, sub-optimal states and investigating RNA 2D conformational equilibria, emerges as a key functional advantage of the BPQ_RNA pipeline.

Our exploration and sampling of multiple sub-optimal configurations aligns with the foundational sub-optimal RNA folding landscape theory established by Wuchty et al. (1999) [33]. Their classical algorithm, RNAsubopt [33, 34], enabled the systematic calculation of all secondary structures within a given energy window, recognizing that functional RNA molecules frequently exist as conformational ensembles rather than resting in a single minimum free energy state. While classical methods algorithmically compute these suboptimal conformers, BPQ_RNA workflow extracts this energetic ensemble directly from the probabilistic quantum measurement profile.

### Future Prospects & Development

The current framework presented in this study bridges experimental thermodynamics and near-term quantum optimization, but several computational and biological avenues remain to be addressed. While BPQ_RNA successfully resolves planar topologies, this initial proof-of-concept currently excludes pseudo-knotted structures. The energetic contributions of pseudoknots are highly dependent on global structural context, making their integration into a strictly two-body quadratic Hamiltonian mathematically challenging without resorting to arbitrary penalization. Given the constraints of low-qubit simulation footprints, our focus remains on validating the thermodynamic mapping of canonical and modified base pairs. Future iterations will explore hybrid classical pre-processing to identify probable pseudoknot interaction regions before quantum encoding, avoiding the pseudoknot overestimation seen in prior models.

By operating at single-base-pair resolution, BPQ_RNA bridges quantum optimization of RNA 2D structure with the thermodynamic consequences of modified RNA. While coarse-grained models scale to longer sequences by treating nucleotide tracts as rigid stems/quartets, BPQ_RNA provides the granularity required to resolve local structural shifts induced by site-specific chemical modifications. This makes the framework inherently integrative: rather than serving solely as a standalone solver, BPQ-RNA might act as a targeted refinement approach within broader computational pipelines, re-evaluating key functionally/therapeutically important motifs where base-pair level resolution is critical to capturing the accurate fold.

While industry-led efforts continue to scale quantum hardware to hundreds of qubits, these large-scale approaches often rely on brute-force mappings that remain highly vulnerable to noise and hardware-specific constraints. Our hybrid pipeline demonstrates that small-scale hybrid approaches, specifically, thermodynamically informed pre-compilation and localized constraint optimization, can help make those utility-scale executions viable. The modular architecture is designed to scale to broader post-transcriptional modification libraries (e.g., pseudouridine and inosine), inherently resolve non-planar pseudoknots, and incorporate higher-order NNDB heuristics, such as terminal mismatches and internal loop asymmetries, to definitively reconcile output free energies with legacy classical algorithms. Finally, as fault-tolerant quantum processing units (QPUs) mature further, we aim to eliminate the classical maturation bottleneck by compiling non-local loop thermodynamics directly into Higher-Order Binary Optimization (HOBO) formulations executed on quantum hardware.

## Data and Code Availability

All code and data required to reproduce the findings of this study are open-source and publicly available. The primary Google Colab execution notebook and integrated workflows are hosted on GitHub at https://github.com/duttan710/BPQ_RNA. The raw execution log and parsed summary data supporting the benchmark (for 20 sequences) presented in this work, are provided in Supplementary Data 1 (Supplementary_Data_1.zip) and also archived alongside the source code in the GitHub repository The text format files for the Nearest Neighbor Database (NNDB)-derived thermodynamic parameters are hosted via GitHub Gist at https://gist.github.com/duttan710/1429a1150dcd7d75fb5ca0cf29b92963/ and linked directly within the main repository and the Colab notebook.

## Supplementary Information

Supplementary table: Table S1. Overview of the results from the adaptive multi-start and multi-replica approach across the 20 test sequences.

**Supplementary Data 1 (Supplementary_Data_1.zip):** Compressed directory containing both the unedited raw QAOA execution logs (results.txt) and the parsed, summary dataset (compiled_qaoa_results.csv) across all 20 benchmark sequences and 5 solver replicas.

## Acknowledgements

We thank the HPE DSI (University of Houston) for the Quantum Computing Course for providing the opportunity to learn about Quantum Computing approaches; Prof. Ansuman Lahiri for constructive feedback on the manuscript; Google Gemini (enterprise-secured Workspace) for coding assistance and language (English) corrections.

## Author contributions

ND, LS, and QV conceptualized the study and analyzed the findings. ND built the workflow, notebook and tutorial. ND wrote the original draft. All authors reviewed and edited the manuscript.

## References

[1] Westhof, E. (2026). Structural Neutrality and Accommodation Shape RNA Evolution. In: Witzany, G. (eds) Integrative Theory of Evolution - Aspects and Insights. Springer, Cham. 10.1007/978-3-032-24295-2_12.

[2] Vicens, Q., & Kieft, J. S. (2022). Thoughts on how to think (and talk) about RNA structure. Proceedings of the National Academy of Sciences of the United States of America, 119(17), e2112677119.

[3] Sacco, G., Bussi, G., & Sanguinetti, G. (2026). Machine learning for RNA secondary structure prediction: a review of current methods and challenges. RNA (New York, N.Y.), 32(4), 443–456.

[4] Westhof, E., Sun, H., Bu, F., & Miao, Z. (2026). The RNA-Puzzles Assessments of RNA-Only Targets in CASP16. Proteins, 94(1), 218–229.

[5] Tinoco, I., Jr, & Bustamante, C. (1999). How RNA folds. Journal of molecular biology, 293(2), 271–281.

[6] Fox, D. M., MacDermaid, C. M., Schreij, A. M. A., Zwierzyna, M., & Walker, R. C. (2022). RNA folding using quantum computers. PLoS computational biology, 18(4), e1010032.

[7] Lewis, M.W., Verma, A. & Eckdahl, T.T. Qfold: a new modeling paradigm for the RNA folding problem. J Heuristics 27, 695–717 (2021).

[8] Jiang, J., Yan, Q., and Li, Y., et al., (2023) Predicting RNA Secondary Structure on Universal Quantum Computer., arXiv: 2305.09561, [quant-ph] https://arxiv.org/abs/2305.09561

[9] T. Zaborniak, J. Giraldo, H. Müller, H. Jabbari and U. Stege, (2022). “A QUBO model of the RNA folding problem optimized by variational hybrid quantum annealing,” 2022 IEEE International Conference on Quantum Computing and Engineering (QCE), Broomfield, CO, USA, pp. 174–185, doi: 10.1109/QCE53715.2022.00037.

[10] Andrews, B., Abraham, J., Fox, D., & Radja, A. (2026). Exploring the Boundaries of Modern Quantum Annealers with RNA Structure Prediction. The journal of physical chemistry. B, 130(9), 2518–2530.

[11] Kumar, V., Alevras, D., Metkar, M., Welling, E., Cade, C., Niesen, I., Friedhoff, T., Park, J.-E., Shivpuje, S., LaDue, M., Davis, W., & Galda, A. (2025). Towards secondary structure prediction of longer mRNA sequences using a quantum-centric optimization scheme. arXiv preprint arXiv:2505.05782 [quant-ph]. 10.48550/arXiv.2505.05782.

[12] Jiang, X., Liu, B., Nie, Z. et al. The role of m6A modification in the biological functions and diseases. Sig Transduct Target Ther 6, 74 (2021). 10.1038/s41392-020-00450-x.

[13] Turner, D. H. & Mathews, D. H. (2009). NNDB: The nearest neighbor parameter database for predicting stability of nucleic acid secondary structure. Nucleic Acids Research. 38, D280–D282.

[14] Mittal, A., Turner, D. H., & Mathews, D. H. (2024). NNDB: An Expanded Database of Nearest Neighbor Parameters for Predicting Stability of Nucleic Acid Secondary Structures. Journal of Molecular Biology, 168549.

[15] Kierzek, E., Zhang, X., Watson, R.M. et al. (2022). Secondary structure prediction for RNA sequences including N<sup><a href=“#ch00fn6”>6</a></sup>-methyladenosine. Nat Commun 13, 1271.

[16] Cerezo, M., Arrasmith, A., Babbush, R. et al. (2021). Variational quantum algorithms. Nat Rev Phys 3, 625–644. (2021).

[17] Farhi, E., Goldstone, J., and Gutmann, S. (2014). A quantum approximate optimization algorithm. arXiv:1411.4028.

[18] Qiskit contributors, “Qiskit: An open-source framework for quantum computing,” 2023.

[19] draw.io Ltd & draw.io AG. (2026). draw.io Desktop (Version 32.5.5) [Computer software]. GitHub. https://github.com/jgraph/drawio-desktop/releases/tag/v32.5.5

[20] Sato, K., Akiyama, M., & Sakakibara, Y. (2021). RNA secondary structure prediction using deep learning with thermodynamic integration. Nature communications, 12(1), 941.

[21] Reuter, J. S., & Mathews, D. H. (2010). RNAstructure: software for RNA secondary structure prediction and analysis. BMC bioinformatics, 11, 129.

[22] Lu, C., Smith, A., Fuchs, R. et al. (2008). Crystal structures of the SAM-III/SMK riboswitch reveal the SAM-dependent translation inhibition mechanism. Nat Struct Mol Biol 15, 1076–1083

[23] Butcher, S., Allain, FT. & Feigon, J. (1999) Solution structure of the loop B domain from the hairpin ribozyme. Nat Struct Mol Biol 6, 212–216

[24] Morosyuk, S. V., Cunningham, P. R., & SantaLucia, J., Jr (2001). Structure and function of the conserved 690 hairpin in Escherichia coli 16 S ribosomal RNA. II. NMR solution structure. Journal of molecular biology, 307(1), 197–211.

[25] Kim, C. H., Kao, C. C., & Tinoco, I., Jr (2000). RNA motifs that determine specificity between a viral replicase and its promoter. Nature structural biology, 7(5), 415–423.

[26] Rozov, A., Demeshkina, N., Khusainov, I., Westhof, E., Yusupov, M., & Yusupova, G. (2016). Novel base-pairing interactions at the tRNA wobble position crucial for accurate reading of the genetic code. Nature communications, 7, 10457.

[27] Jeong, H., & Kim, J. (2021). Unique anticodon loop conformation with the flipped-out wobble nucleotide in the crystal structure of unbound tRNAVal. RNA (New York, N.Y.), 27(11), 1330–1338.

[28] Zhou, J., Lancaster, L., Donohue, J. P., & Noller, H. F. (2014). How the ribosome hands the A-site tRNA to the P site during EF-G–catalyzed translocation. Science. https://doi.org/1255030

[29] Duszczyk, M. M., Wutz, A., Rybin, V., & Sattler, M. (2011). The Xist RNA A-repeat comprises a novel AUCG tetraloop fold and a platform for multimerization. RNA (New York, N.Y.), 17(11), 1973–1982.

[30] Jones, A. N., Tikhaia, E., Mourão, A., & Sattler, M. (2022). Structural effects of m6A modification of the Xist A-repeat AUCG tetraloop and its recognition by YTHDC1. Nucleic Acids Research, 50(4), 2350–2362.

[31] Yang, S., Pham, N. T., Li, Z., et. al., (2025). Advances in RNA secondary structure prediction and RNA modifications: Methods, data, and applications. arXiv preprint arXiv:2501.04056. 10.48550/arXiv.2501.04056.

[32] Alevras, D., Metkar, M., Yamamoto, T., et al., “mrna secondary structure prediction using utility-scale quantum computers,” in 2024 IEEE International Conference on Quantum Computing and Engineering (QCE), vol. 1. IEEE, 2024, pp. 488–499.

[33] Wuchty, S., Fontana, W., Hofacker, I. L., and Schuster P. (1999), “Complete Suboptimal Folding of RNA and the Stability of Secondary Structures”, Biopolymers: 49,145–165

[34] Lorenz, R., Bernhart, S.H., Hoener zu Siederdissen, C. et. al. (2011), “ViennaRNA Package 2.0”, Algorithms for Molecular Biology: 6:26

