## Supplementary Information for "BPQ_RNA: A Thermodynamically Informed, Hybrid Quantum-Classical Framework for RNA 2D Structure Prediction Including Chemical Modifications"

**Table S1:** Overview of the results from the adaptive multi-start and multi-replica approach across the 20 test sequences

| Sequence | Length (nt) | Qubits | Core stack energy (kcal/mol) | MFE State Recovery (out of 5) | Average number of Cycles till Convergence | Replicas Triggering Early Stop |
|---|---|---|---|---|---|---|
| **S1** | 10 | 7 | -5.83 | 5 / 5 | 1.4 | 5 / 5 |
| **S2** | 11 | 5 | -3.32 | 5 / 5 | 1.0 | 5 / 5 |
| **S3** | 12 | 16 | -7.64 | 5 / 5 | 12.6 | 5 / 5 |
| **S4** | 12 | 16 | -6.54 | 3 / 5 | 10.8 | 3 / 5 |
| **S5** | 13 | 8 | -9.14 | 5 / 5 | 1.2 | 5 / 5 |
| **S6** | 13 | 14 | -6.44 | 5 / 5 | 6.2 | 5 / 5 |
| **S7** | 13 | 15 | -5.83 | 5 / 5 | 4.6 | 5 / 5 |
| **S8** | 13 | 17 | -6.14 | 5 / 5 | 11.2 | 5 / 5 |
| **S9** | 13 | 18 | -11.05 | 4 / 5 | 12.0 | 4 / 5 |
| **S10** | 13 | 18 | -4.54 | 5 / 5 | 7.8 | 4 / 5 |
| **S11** | 14 | 12 | -6.74 | 5 / 5 | 2.6 | 5 / 5 |
| **S12** | 14 | 12 | -6.34 | 5 / 5 | 3.8 | 5 / 5 |
| **S13** | 14 | 16 | -4.54 | 4 / 5 | 8.0 | 4 / 5 |
| **S14** | 15 | 13 | -6.74 | 5 / 5 | 2.4 | 5 / 5 |
| **S15** | 16 | 14 | -6.74 | 5 / 5 | 2.6 | 5 / 5 |
| **S16** | 11 | 14 | -5.43 | 5 / 5 | 7.0 | 5 / 5 |
| **S17** | 11 | 14 | -5.43 | 5 / 5 | 13.6 | 5 / 5 |
| **S18** | 12 | 16 | -9.24 | 3 / 5 | 12.8 | 3 / 5 |
| **S19** | 12 | 16 | -9.24 | 4 / 5 | 15.0 | 4 / 5 |
| **S20** | 13 | 22 | -5.74 | 1 / 5 | 18.8* | 1 / 5 |

**Note for sequence S20:* Replicates 1, 2, 3, and 5 ran to the maximum 20-cycle limit, while Replicate 4 triggered early stopping at Cycle 14 upon finding the global minimum (average 18.8 attempted cycles, 12.8 executed cycles excluding skipped cycles with sampling error).

Complete cycle-by-cycle solver outputs, raw quantum objective values, intermediate energy trap evaluations, and full top-$K = 5$ sampled bitstrings for each replicate are provided as a standalone plain-text file in **Supplementary Data 1** (results.txt).
